# Low-pH conditions drive transient changes in shell calcification and the microbiome in a pH-resistant strain of the the Pacific oyster *Magallana gigas*

**DOI:** 10.1101/2025.04.13.648625

**Authors:** Roxanne M.W. Banker, John J. Stachowicz, David A. Gold

**Affiliations:** Department of Biology, Providence College, Rhode Island, United States of America; Department of Earth and Planetary Sciences, University of California, Davis, California, United States of America; Department of Evolution and Ecology, University of California, Davis, California, United States of America; Bodega Marine Laboratory, University of California, Davis, California, United States of America

**Keywords:** ocean acidification, oysters, microbiome, gene expression

## Abstract

The study explores the effects of elevated pCO_2_ on shell calcification, microbiome composition, and gene expression in a strain of Pacific oyster (*Magallana gigas*) selectively bred for low-pH resistance. Juvenile oysters reared under low-pH conditions exhibited increased shell mass compared to the control population by 51 days post-fertilization, despite high variance in shell size at earlier stages. Microbiome analyses revealed significant shifts in community composition under low-pH conditions, particularly in bacterial taxa involved in CO_2_ production and biogeochemical cycling, which could influence carbonate chemistry within oyster tissues. Gene expression profiling demonstrates differential regulation of genes related to biomineralization, immunity, and microbial interactions under low-pH conditions. For example, multiple carbonic anhydrases exhibited treatment-specific expression patterns, suggesting a role in adapting to low-pH environments. Observed changes in immune-related genes imply a relaxation of immune responses, potentially reflecting resource reallocation toward calcification processes. These results collectively support the “dysbiosis hypothesis,” where oysters adapt to environmental stress by modulating their microbiomes and gene expression. Future studies should investigate whether these responses are consistent across oyster strains and environmental conditions, providing insights into the resilience of aquaculture species to ocean acidification.

**SUMMARY STATEMENT:** Elevated pCO_2_ impacts Pacific oyster calcification, microbiome composition, and gene expression, suggesting genetic adaptation and microbiome shifts may be crucial for resilience to ocean acidification.

## INTRODUCTION

The impact of ocean acidification on marine life is an area of great concern and uncertainty. Ocean acidification occurs when a rapid increase of atmospheric carbon dioxide causes seawater to absorb CO₂ faster than the ocean’s natural buffering system can counteract, leading to a drop in pH [1–3]. In principle, this is particularly problematic for marine species that builds calcium carbonate skeletons–such as bivalves and corals–because lower seawater pH potentially limits availability of the materials required to precipitate a calcium carbonate (i.e. [CO ^2-^]) skeleton [3–7]. Despite these dire implications, marine species with calcium carbonate skeletons show unpredictable responses to seawater acidification. Some species appear particularly sensitive, while others demonstrate resilience [8–10]. One possible explanation for this resiliency is the buffering role resident microbes may play [11,12], but research supporting this hypothesis is currently scarce.

The Pacific oyster *Magallana gigas* (formerly known as *Crassostrea gigas*) is an emerging model system to study the interplay between host and microbes in shell formation [13–15]. Like many intertidal organisms, *M. gigas* is a resilient species, able to withstand large daily fluxes in temperature and salinity. Its shell growth is highly plastic, thanks in part to its unusual ability to precipitate chalk within its shell [16]. There is some evidence that microbial community composition in *M. gigas* correlates with changes in calcification and microstructural morphology in the shell [11]. For example, chemical manipulation of the oyster microbiome results in temporary changes in shell growth and microbiome composition, but these differences are lost over time [17]. The return of a normal microbiome and shell growth rates correlate with the expression of oyster genes that are involved in host-microbe interactions, suggesting the animal plays an active role in regulating its microbiome alongside shell formation. Others have examined how oysters and related bivalves specifically respond to acidification, but the results have been mixed. For example, different studies have found that lowering seawater pH can increase or decrease oyster shell growth ([18], [19], [20], [21]). Similarly, there is evidence that bacterial richness is altered under low-pH conditions ([22][23]), but the details of these processes vary, and require further exploration [24].

Here we use juveniles of *M. gigas* to better characterize how oyster gene expression, microbial community composition, and shell formation respond to changes in seawater pH during a critical early life history stage. We reared oysters from larvae to the juvenile stage under two conditions: a control pH (∼8) and low pH (∼7.7) that provides a simulation of near-future ocean acidification scenarios [25]. Notably, animals obtained for this experiment were bred by the Molluscan Broodstock Program at Oregon State University in order to be resistant to low pH conditions. Thus, results from this work have important implications for how organisms bred to resist the deleterious effects of global change will respond to declining marine conditions. Collectively, these data allow us to disentangle animal-versus microbiome-driven effects of calcification, and how these factors might respond to changing ocean conditions.

## MATERIALS AND METHODS

The results of this study are part of a larger experiment first described in Banker et al. (2022). We will redescribe relevant portions of the experiment here, as well as include additional information related to pH manipulation, which was not considered in the original study.

### Study organism and broodstock treatment

We obtained broodstock oysters from the Hog Island Oyster Company (HIOC) on July 30, 2019. Broodstocks were rinsed and scrubbed with freshwater upon arriving at the Bodega Marine Laboratory (BML) to remove epibionts, and were then soaked in a 60 ppm solution of sodium hypochlorite (bleach) for 1-hour according to regulations from the California Department of Fish and Wildlife. Oysters were left to dry in air for ∼1.5 hours before being placed in filtered seawater at approximately 18℃. Broodstock were fed daily with 5 gallons of algal culture composed of either *Isochrysis sp.* (strain CCMP463, Bigelow National Center for Marine Algae and Microbiota) or *Nannochloropsis sp.* (strain CCMP525, Bigelow National Center for Marine Algae and Microbiota). On days when cultures crashed or were not viable, algal paste (Reed Mariculture, Shellfish Diet 1800) was used (Table S1). Water was flushed every 2-3 days by inserting a standing pipe into the drain at the bottom of the tank and allowing incoming water to displace tank water. This was done for 30 minutes for each flush. A full water change, including a complete draining and cleaning of the tank, occurred every 7 days.

### Spawning oysters

We obtained gametes from reproductive adults, produced fertilized embryos, and developed larvae using the methods in Banker et al. (2022). After larvae developed into the D-hinge veliger stage, they were separated evenly into 9 buckets (three treatments, three bucket replicates each) at a density of approximately 10 larvae/mL (Table S1).

### Experimental design

Oysters in all treatments were maintained in 10-gallon food-grade buckets. Low-pH conditions were maintained at a pH of ∼7.70 using a Pinpoint^®^ pH controller coupled with the Pinpoint^®^ pH regulator (American Marine, Inc.) connected to a tank of compressed air composed of 1% (10,000 ppm) CO_2_ (Airgas). When pH in the low-pH treatment buckets increased above a predetermined set point (pH=7.80), the controller would activate the solenoid in the regulator to input CO_2_ into the system, thus driving down pH. All treatments were supplied with air through an airstone to maintain oxygen in the tanks.

Crushed oyster shell, size sorted to 175 um and autoclaved, was added to buckets on September 16 (20 days post-fertilization) to act as a substrate for settlement (Table S1). Daily observations of buckets indicated that settlement of oysters began on day 26 and the remainder of individuals settled by day 31 post-fertilization (Table S1). Despite the addition of crushed shell material, juvenile oysters settled exclusively on the interior surfaces of buckets. Juvenile oysters at this stage were collected by scraping a pea-sized volume of settled juvenile oysters off of the sides of buckets using a small spatula. Oyster samples were quickly rinsed in ethanol to remove any external microbes before being frozen at -80 °C. On Day 38 post-fertilization, two of the three replicate buckets from both treatments were destructively sampled to collect juvenile oysters for nucleotide extraction at an additional time point during ontogeny.

The naming scheme of the samples is “Treatment-Time-Bucket-Replicate”. For example, the three replicates from control bucket 1 are referred to as “C-38D-1-[1–3]” and the three replicates from bucket 2 of the low-pH treatment are “LP-38D-2-[1–3]”. Three replicate buckets were maintained for each treatment in case a mass mortality event affected one or more buckets. When all experimental buckets made it past settlement without issue, we destructively sampled two of the three buckets. We grew out the final replicate bucket of each treatment to 51 days post-settlement to capture an additional ontogenetic time point of oyster development with our sampling and analyses. However, the lack of biological replicates for 51-day samples limited the types of analyses we could perform on these data.

### Oyster Maintenance

Oyster D-hinge veligers in buckets were fed a diet of *Isochrysis sp.* at approximately 30,000 cells/mL from the time larvae were transferred to buckets until September 4th (8 days post-fertilization). Subsequently feeding concentrations were gradually increased according to established animal care protocols used by the Aquatic Resources Group (ARG) at BML. Algal feed concentrations in oyster tanks were raised to ∼100,000 cells/mL by September 19 (23 days post-fertilization). When settlement was first observed in buckets on September 22 (26 days post-fertilization), feeding was increased to *ad libitum* to encourage growth.

To maintain the culture quality of the static system, water in buckets was manually changed every three days. For the first water change that took place on September 1 (5 days post-fertilization) a 35 um sieve was used to isolate larvae. Progressively larger sieves were used over the course of the experiment as oysters grew to larger sizes (Table S1).

After water changes, water used to restock oysters for the acidified treatment was taken from incoming filtered seawater, which had a pH of approximately 7.8, but was not otherwise pretreated. For the Control treatment, water was pretreated by adding incoming seawater to a separate tank, and bubbling it with an airstone to raise the pH to ∼8.0.

### Water Chemistry Monitoring

Temperature, pH, dissolved oxygen (DO), and salinity were recorded twice daily for each experimental tank to ensure that habitable conditions were maintained in all microcosms. This was done using a Pinpoint^®^ pH controller and probe, a Sper Scientific Dissolved Oxygen Meter Kit (which measured both DO and temperature), and a refractometer. In addition to daily monitoring, oyster culture water was sampled during each water change to gain a more precise measurement of seawater pH conditions. Water samples were also taken at this time from the source of filtered seawater used in this experiment, starting on the water change that occurred on September 7, 2019.

Each water sample was analyzed spectrophotometrically for pH. Samples were collected in 125 mL glass bottles with a positive meniscus, and then spiked with mercuric chloride. Caps were wrapped with parafilm and bottles were stored in 4°C until analysis. These samples were analyzed on an Ocean Optics Jaz Spectrophotometer EL200 (SD +/- 0.003) using *m*-cresol purple [26]. A calibration regression was produced for each batch of dye (*m-*cresol) and calibrated against Tris for a <0.1 pH offset in order to correct measurements.

Samples for total alkalinity were collected in 250 mL Nalgene© tubes with a small amount of headspace, and immediately frozen at -20 °C. These were run on an automated Gran titration on a Metrohm 809 Titrando (SD +/- 4.2 umol/kg); acid concentrations standardized using Dickson certified reference materials.

Mean alkalinity was calculated for each bucket on the basis of samples that were successfully analyzed for that bucket, though the number of samples that was used for each alkalinity bucket mean was variable (S2 Table). Mean total alkalinity values for each bucket were assigned to each spectrophotometry derived pH value that also belonged to the same bucket.

These average alkalinity values for each bucket were used as a final correction to pH measurements, using CO2calc [27] with pK_1_ and pK_2_ CO_2_ equilibrium constants from [28] and KHSO_4_ from [29]. All uncorrected and corrected pH data can be found in Table S2.

To confirm that the experimental design was effective in regards to maintaining different pH conditions, a mixed effects model was used to compare mean pH among treatment groups while accounting for bucket ID as the random effect. This test was performed in R v4.3.1 using the lmer package as follows: lmer(pH.corrected ∼ treatment + (1 | bucket/treatment). This procedure was performed with pH values measured on the Ocean Optics Jaz Spectrophotometer.

### 16s DNA Extraction, PCR Amplification, and Sequencing

ZymoBIOMICS DNA Miniprep Kits were used to extract total DNA from larval (1000s individuals) and juvenile (100s of individuals) samples. All samples were taken in duplicate so that one sample of each pair was treated to extract DNA while the other was treated to extract RNA. For DNA extraction, test kits were also applied to three empty microcentrifuge collection tubes (those that had been used to collect samples) and two bead tubes (from the extraction kit) as negative controls to assess for possible sources of contamination.

Total DNA was sent to the Integrated Microbiome Resource (IMR; https://imr.bio/) at Dalhousie University (Halifax, Nova Scotia), where PCR amplification and sequencing of samples was carried out. Sequencing libraries were prepared at by targeting the V4-V5 regions of the 16s rRNA gene for amplification using the the 515F (GTGYCAGCMGCCGCGGTAA) and 926R (CCGYCAATTYMTTTRAGTTT) primer set. Sequencing was performed on the Illumina HiSeq platform.

### Microbiome Analysis and Visualization

The resulting fastq files were analyzed in R (v. 4.3.1) using dada2 (v. 1.28.0), phyloseq (v. 1.44.0), vegan (v. 2.6.4), FSA (v. 0.9.4), and ggplot2 (v. 3.4.2) (Dixon, 2003; Wickham, 2009; McMurdie & Holmes, 2013; Callahan et al., 2016; Ogle, 2016; R Core Team, 2016; Davis et al., 2018). A detailed walkthrough of subsequent analyses in R can be found in R-markdown summary file hosted on github (https://github.com/Roxanne-Banker/Oyster_OA).

Reads were truncated to reduce estimated rates of error and optimize sequence merging in subsequent steps using the filterAndTrim function in dada2. The dada command in the same package was used to denoise forward and reverse reads; these reads were then merged using mergePairs. These steps produced a table of amplicon sequence variants (ASVs) and chimeric sequences were removed with the removeBimeraDenovo command. Taxonomy was assigned using version 132 of the SILVA taxonomic training set from DADA2 [30–32].

The resulting ASV and taxonomy tables were passed as a phyloseq object for subsequent decontamination and analyses. The prevalence method in Decontam was employed to identify contaminants in negative control samples using a threshold of 0.1. This process indicated 1 possible contaminating sequence, which was removed from the dataset. Chloroplast, mitochondria, and animal sequences were additionally filtered out.

### Alpha Diversity

Within sample diversity, i.e., alpha diversity, was estimated using three diversity metrics: observed ASVs and the Shannon diversity index using the estimate_richness function in phyloseq. The Kruskal-Wallis test was used to compare whether buckets differed in terms of any of these diversity metrics.

### Ordination

Sequence counts in each sample (Table S3) were subsequently transformed using varianceStabilizingTransformation in DESeq2 [33]. Negative values that resulted from this log-transformation method were replaced with zeroes to enable ordination. Ordination was performed by transforming sequence counts into relative abundances and calculating Bray-Curtis distances from those data. The ordinate function from phyloseq was used using the non-metric multidimensional scaling (NMDS) method.

### Beta diversity

A PERMANOVA - permutational analysis of variance- was used to assess statistical significances of among group differences in composition (i.e., beta diversity) [34]. The PERMANOVA was applied to Bray-Curtis distances, using 9,999 permutations to account for multiple tests, with the adonis2 function in vegan. The PERMANOVA test is less sensitive than other multivariate tests to multiple comparisons [35], results may be affected by differences in variation (i.e., dispersion) among groups [36]. Differences of dispersion among sample categories was assessed with the betadisper function in vegan, using 999 permutations using permutest.

As performing multiple comparisons is likely to increase the rate of Type I error, p-values resulting from the PERMANOVA and Levene’s tests were adjusted using the Bonferroni adjustment. This was implemented in the p.adjust function in R. A critical threshold of α = 0.05 was used to assess significance for all tests.

### Community Composition Analyses

To assess and compare bacterial community composition, ASVs were collapsed to the family level using the tax_glom function in phyloseq and taxa that varied less than 0.20% in abundance between samples were removed. This threshold was selected to remove taxa that did not vary between samples, and to remove rare taxa that might produce false positives during subsequent comparative tests. The average abundance of families were compared between treatment groups, and also within treatment groups by age, using the Kruskal-Wallis test at a critical threshold of 0.05. When the Kruskal-Wallis test was rejected, the Dunn test was applied as a post-hoc, and Bonferroni correct p-values were used to assess which pairwise comparisons for a given family were significant (α = 0.05).

Phylogenetic Investigation of Communities by Reconstruction of Unobserved States (PICRUSt2), software used to predict functional abundances based on 16s rRNA sequences was applied to our data [37–42]. The full PICRUST2 pipeline was applied to 16s counts prior to the variance stabilizing transformation in DESeq2. The command to execute the program pipeline was: *picrust2_pipeline.py -s study_seqs.fna* -*i study_seqs.biom -o picrust2_out_pipeline -p 1 -- metagenome_contrib*. The output of this command predicted KEGG (Kyoto Encyclopedia of Genes and Genomes) pathways, or the functional potential of the community of microorganisms based on the phylogenetic profiles of the 16S rRNA gene sequences collected during our experiment. This is done by associating the sequences with specific KEGG Orthologs (KOs), which are linked to known metabolic and cellular pathways in the KEGG database. KO abundances were read into R where the package ggpicrust2 was used to calculate differential abundances of orthologs using the DESeq2 protocol. P-values for assessing significance of the differential abundance analysis were corrected for multiple comparisons using the Bonferroni correction. We further investigated orthologs that showed a significant difference (corrected p-value < 0.05) in predicted expression between C38 versus OA38. Moreover, we focused on orthologs that belong to biochemical reactions (KEGG Modules) that are known to be involved with carbonate chemistry in free living microbial mats [43], following the approach applied by [44]. Microbial metabolic reactions that are known to increase alkalinity and carbonate saturation state that we investigated include denitrification, nitrate reduction, sulfate reduction, acetoclastic methanogenesis, hydrogenotrophic methanogenesis, photosynthesis, and photoautotrophy. By contrast, microbially driven reactions that are known to decrease carbonate saturation state include sulfide oxidation, methylotrophic methanogenesis, and ammonium oxidation. We used ggplot2 in R to visualize the number of orthologs belonging to each pathway and the percent contribution of microbial phyla to these pathways.

### Oyster RNA extraction, sequencing, and bioinformatics

Total RNA was isolated from oyster tissues using 0.50 mL of TRIZOL Reagent (Life Technologies). The tissue was homogenized in TriZol using a Pellet Pestle^®^ Motor (Kontes). The samples incubated at room temperature for 5 minutes before adding 0.10 mL of chloroform.After another ∼2 minutes of incubation at room temperature, the samples were centrifuged at 12,000xg at 4°C for 15 minutes. The aqueous layer was transferred to a new tube and combined with an equal volume of isopropyl alcohol and 1 μg of glycol blue. After a 10 minute incubation at room temperature, the samples were centrifuged a second time at 12,000xg at 4°C for 15 minutes. The resulting RNA pellet was washed in 70% ethanol and then dried for ∼5 minutes. Finally, the pellets were dissolved in ∼40μl of RNase free water. RNA quality was determined using a NanoDrop (NanoDrop Technologies, Inc.). The samples were submitted to the UC Davis DNA Technologies & Expression Analysis Core Laboratory, for cDNA preparation 3’-TAG sequencing.

Detailed code for the bioinformatics is provided on GitHub (https://github.com/DavidGoldLab/2025_Oyster_RNA-Seq). Briefly, the reads were cleaned with BBMAP v39.01, and then mapped to the *C.gigas* "xbMagGiga1.1" genome assembly (NCBI accession GCF_963853765.) with STAR v2.7.6a. We then created an updated set of gene models and tested for differential gene expression using the Cufflinks v.2.2.1 package. We used the PtR program packaged with Trinity v2.11.0 to generate heat maps and PCA plots of the data. Enrichment analyses were performed on the NIH DAVID v2024q2 server [45] using “OFFICIAL_GENE_SYMBOL” identifiers, “*Crassostrea gigas*” as the species, and a list of all differentially expressed genes as the “Background”.

### Scanning Electron Microscopy (SEM) and Shell Analysis

On Days 38 and 51, additional oyster samples were removed from their affixed position on buckets by carefully removing them with a paintbrush. These oysters were stored in ethanol until they could be processed. Oyster shells were dried and mounted onto an aluminum stub using carbon tape and then viewed under a Hitachi TM300 scanning electron microscope (SEM). This allowed for clear visualization of the prodissoconch II (pre-settlement larval shell) and the dissoconch (post-settlement juvenile shell) to calculate different shell growth metrics.

The visual analysis program ImageJ was used to measure the areas of the larval and juvenile shells of specimens from each treatment bucket. A linear mixed effects model was fitted using restricted maximum likelihood in the lme4 package in R to assess between treatment differences in shell size and if there was a bucket effect on shell growth [46,47]. The model predicted shell area and mass, respectively, with treatment as a fixed-effect parameter and bucket ID as the random-effect: lmer(area.um2 ∼ treatment + (1 | bucket/treatment). The mixed effects models were only applied to samples collected 38 days post-fertilization because there were no bucket replicates collected 51 days post-fertilization. Ten separate shells from each bucket were also weighed using a Sartorius Pro 11 digital scale to collect shell mass data in addition to shell area. The same statistical procedure was applied to shell mass as the shell area data. Because there were bucket replicate samples collected at 38 days but not 51 days post fertilization, we assessed differences in shell size (area and mass) at 51 days using a t-test instead of the mixed linear model that we applied to 38 day oysters in order to tease apart shell differences driven by bucket identity versus treatment. These t tests were only applied to 51 day bucket samples, which are pseudoreplicates. Three pseudoreplicates were taken from the control bucket and the low-pH bucket.

Average percent difference of oyster shell area and mass between the control and low-pH treatments was calculated by first taking the mean shell area of 38 day old shells from both treatments. The percent difference was then calculated as (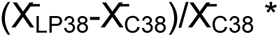 * 100, where 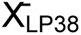 is the mean shell area (mm^2^) for 38 day old low-pH oysters and 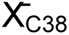 is the mean shell area (mm^2^) for 38 day old control oysters. This calculation was also used to assess percent area and mass differences between control and low-pH oysters at 51 days.

## RESULTS

### Oyster growth

Temperature, salinity, and dissolved oxygen were statistically indistinguishable between treatments for the duration of the experimental period, whereas pH differed significantly between the control and low-pH conditions (Table 1). In terms of shell area, low-pH oysters were on average 136% larger than control oysters at Day 38, but only 45% larger than control oysters by Day 51 (Fig. 1). Similarly, low-pH oysters were 38% heavier than control oysters at 38 days, and 186% heavier at 51 days (Fig. 1). However a strong interaction between treatment and bucket ID complicates the interpretation of these results. Approximately 52% of the variance of shell area can be accounted for by the interaction between treatment and bucket ID; in contrast, none of the variance of shell mass was attributed to the treatment (Tables 2 & 3). Follow-up pairwise F-tests on shell area revealed that variance is statistically different in multiple comparisons between control buckets and low-pH buckets at 38 days. Significant pairwise F-test comparisons for shell area are listed in Table 4, though all pairwise tests among bucket samples collected at 38 days were performed. This result indicates that the low-pH oysters generally display greater variability in shell area than their control counterparts.

**Figure 1.**
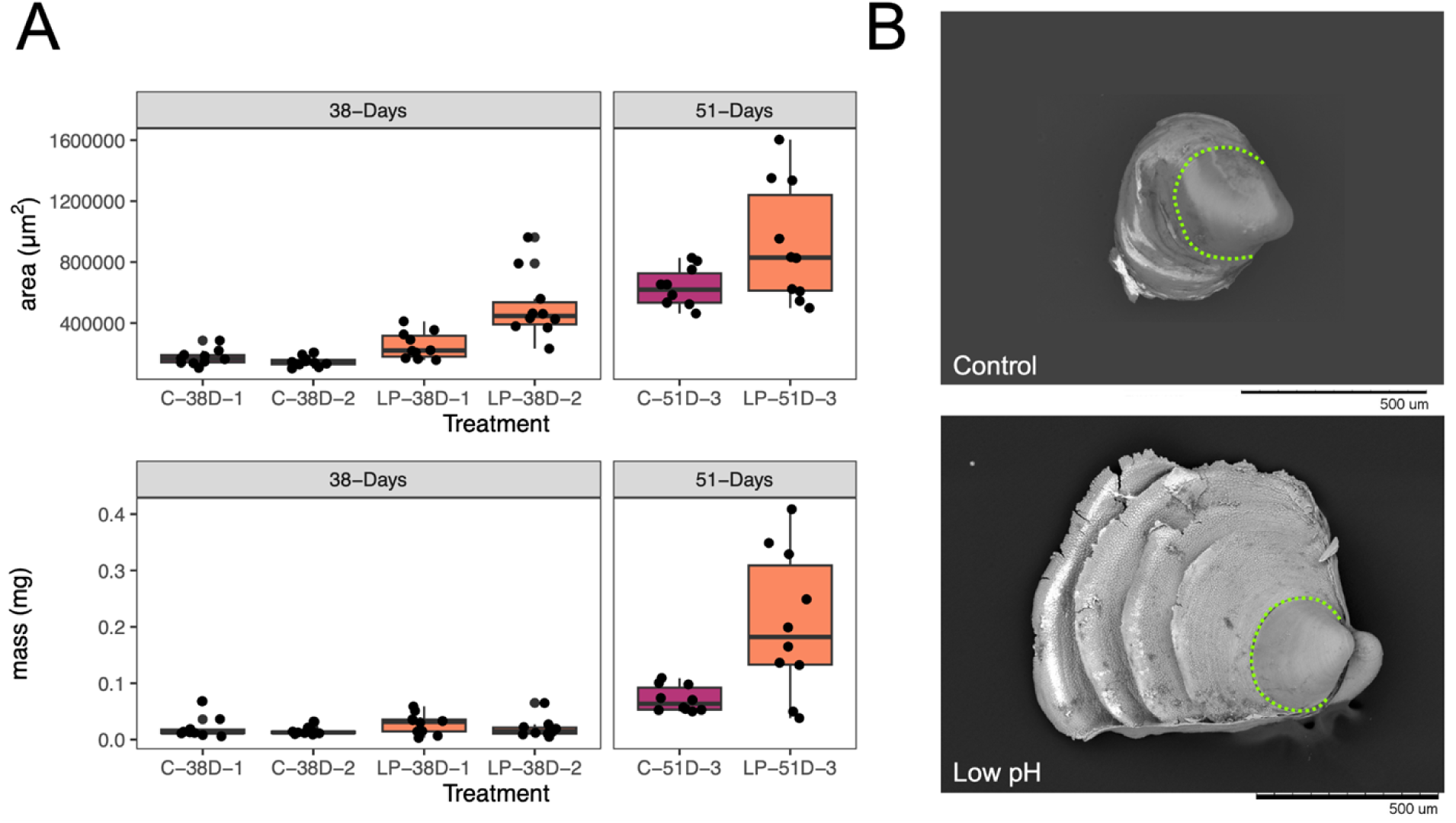
Changes in shell area and mass in oysters over 51 days. (A) Boxplots of whole shell area (top) and mass (bottom) for control and low-ph oysters collected at 38 days post-fertilization (left) and 51 days post-fertilization (right). N=10 for each bucket. (B) Example SEM images showing the variation in shell growth between individual oysters. The larval shell boundary is noted with a green dotted line.

**Table 1.** Summary of the linear mixed effects model fit to pH as the outcome variable.

**Fixed effects (treatment):**
| Effect | Estimate | Std. Error | t-value | p-value |
| --- | --- | --- | --- | --- |
| Intercept<br>(control) | 7.94480 | 0.02067 | 384.43 | 2.3•10<sup>-9*</sup> |
| low-pH | -0.29881 | 0.02923 | -10.22 | 0.000908* |

**Random Effects:**
| Grouping | Effect | Variance | % Variance | Std. Dev |
| --- | --- | --- | --- | --- |
| bucket ID:treatment | Intercept | $8.679 \bullet 10^{-05}$ | 0.57 | 0.0093161 |
| bucket ID | Intercept | $4.098 \bullet 10^{-07}$ | 0.01 | 0.0006402 |
| Residual | | $1.512 \bullet 10^{-02}$ | 99.42 | 0.1229619 |

**Table 2.** Summary of fixed and random effects from linear mixed effects model fit to log(<u>whole shell area (µm^2^)</u>) as the outcome variable for 38 day-old oysters only.

**Fixed effects (treatment):**
| Effect | Estimate | Std. Error | t-value | p-value |
| --- | --- | --- | --- | --- |
| Intercept<br>(control) | 11.9554 | 0.2466 | 48.488 | 0.000425* |
| low-pH | 0.7688 | 0.3487 | 2.205 | 0.158281 |

**Random Effects:**
| Grouping | Effect | Variance | %<br>Variance | Std. Dev |
| --- | --- | --- | --- | --- |
| bucket ID:treatment | Intercept | 0.1110313 | 52.4471 | 0.33321 |
| bucket ID | Intercept | 0.0005462 | 0.2580048 | 0.02337 |
| Residual |  | 0.1001240 | 47.29489 | 0.31642 |

**Table 3.** Summary of fixed and random effects from linear mixed effects model fit to log(<u>shell mass (mg))</u> as the outcome variable for 38 day-old juvenile oysters only.

**Fixed effects (treatment):**
| Effect | Estimate | Std. Error | t-value | p-value |
| --- | --- | --- | --- | --- |
| **Intercept**<br>**(control)** | -1.84019 | 0.06806 | -27.039 | $2.0 \bullet 10^{-16} *$ |
| **low-pH** | 0.09651 | 0.09625 | 1.003 | 0.322 |

**Random Effects:**
| Grouping | Effect | Variance | % Variance | Std. Dev |
| --- | --- | --- | --- | --- |
| **bucket ID:treatment** | **Intercept** | 0.000000 | 0.00 | 0.00000 |
| **bucket ID** | **Intercept** | 0.000000 | 0.00 | 0.00000 |
| **Residual** |  | 0.09263 | 100.00 | 0.3044 |

**Table 4.** F-test comparison to assess bucket variance of shell area. (*) means p < 0.05. Bonferroni adjusted p-value (p_B_), k=6.

| Comparison | p-value | $p_B$ |
| --- | --- | --- |
| C-38D-1 vs. C-38D-2 | $2.230294 \bullet 10^{-01}$ | 1 |
| C-38D-1 vs. LP-38D-1 | $1.108396 \bullet 10^{-01}$ | $6.650376 \bullet 10^{-01}$ |
| C-38D-1 vs. LP-38D-2 | $1.904504 \bullet 10^{-04}$ | $1.142702 \bullet 10^{-03} *$ |
| C-38D-2 vs. LP-38D-1 | $7.294989 \bullet 10^{-03}$ | $4.376994 \bullet 10^{-02} *$ |
| C-38D-2 vs. LP-38D-2 | $5.255807 \bullet 10^{-06}$ | $3.153484 \bullet 10^{-05} *$ |
| LP-38-D1 vs. LP-38D-2 | 1.395662●10<sup>-02</sup> | 8.373969●10<sup>-02</sup> |

A t-test was applied to 51 day shell metrics because we lacked bucket replicates at this stage to apply a mixed linear model. The difference in shell mass at 51 days between low-pH and control oysters is statistically significant (T-test; t(11.559)= 3.1905, p < 0.05). This t-test was applied to pseudoreplicate samples taken from one bucket for each treatment at 51 days (n=10 shells for each bucket, see Materials and Methods for explanation of uneven sampling design).

In summary, oysters raised under low-pH conditions exhibited a higher range of variation in shell area at 38 days. By 51 days, these oysters’ shells were significantly more massive than those raised under normal (pH=8) conditions, with the caveat that we only have one independent replicate of each treatment for day 51 and buckets varied in their treatment effect at day 38.

### Microbial community composition

While there was no difference in microbe species diversity (Kruskal-Wallis: Observed ASVs: χ^2^(5) = 5.4678, p > 0.05; Shannon diversity index: χ^2^(5) = 10.029, p > 0.05; Fig. 2) between treatments, there were major shifts in community composition based on oyster age and pH.

**Figure 2.**
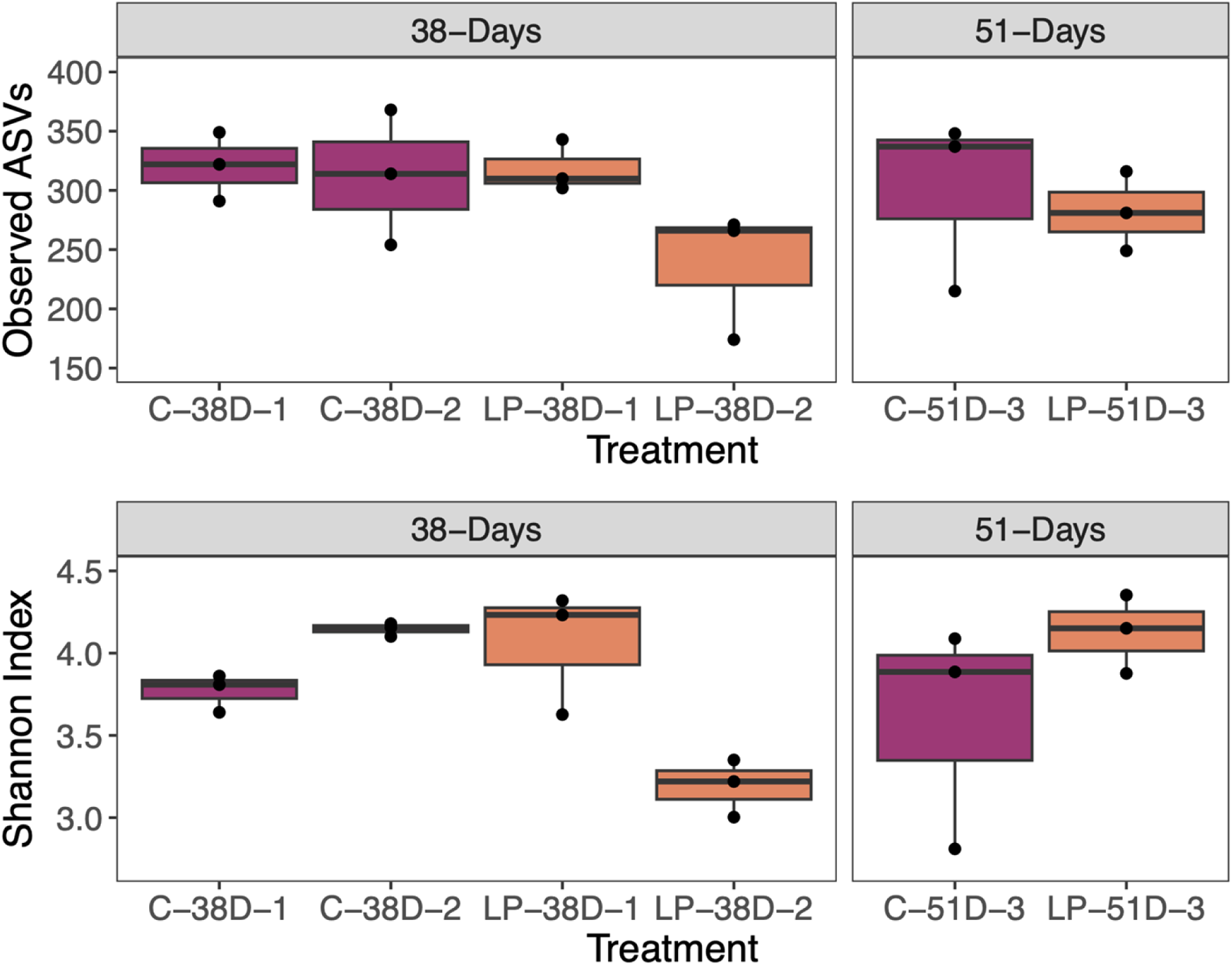
Changes in microbiome composition in oysters over 51 days. Observed amplicon sequence variants (ASVs) and Shannon index were calculated for each sample. C: Control; LP: Low-pH. N=3 for each bucket.

Beta diversity differed when comparing oysters based on treatment identity as well as when grouped based on treatment identity and age (Table 5). Permanova results show that for pairwise comparisons among treatment-time groups, only the comparison between samples at 51 days (C-51D-3 versus LP-51D-3) was not significant (Table 6). The comparison among treatment-time groups also yielded a significant heterogeneity of dispersions test, indicating that the different treatment-time groups have unequal variances (Table 5). Pairwise comparisons show that LP-38D and LP-51D buckets have significantly different variances, again supporting the hypothesis that low-pH oysters exhibited high degrees of variance at 38 days (Table 6; Fig. 3). Thus, while samples from all control buckets were statistically distinct from low-pH buckets at 38 days, oysters had statistically similar microbiomes by Day 51 post fertilization (Table 6; Fig. 3).

**Figure 3.**
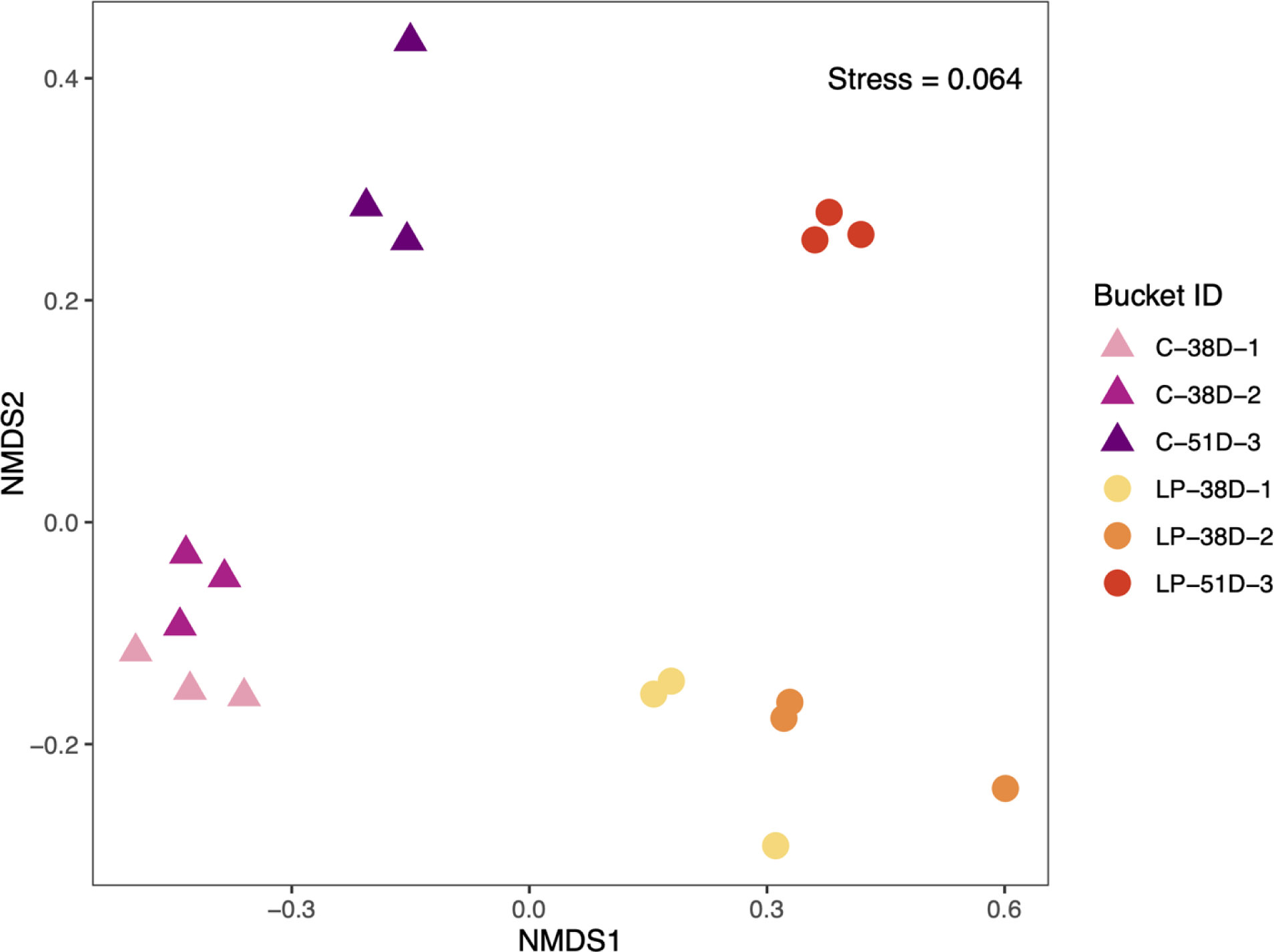
Non-metric multidimensional scaling visualization based on Bray-Curtis dissimilarities of bacterial communities. Resultant stress of the NMDS ordination was 0.064, indicating that the 2-D NMDS plot is a good representation of Bray-Curtis distances among samples [48].

**Table 5.** Summary of results of PERMANOVA and Levene’s test for homogeneity of dispersions for tests comparing Control and Low-pH treatments, and all time points within each treatment. A significant p-value for PERMANOVA indicates that the groups tested are statistically distinct. A significant p-value for Levene’s test indicates that the group dispersions of the groups tests are statistically different, which breaks one of the assumptions of PERMANOVA. Bonferroni adjusted p-values (p_B_) have been corrected for k=2 tests, and (*) indicates p < 0.05.

|  | PERMANOVA |  |  |  | Levene's test |  |  |
| --- | --- | --- | --- | --- | --- | --- | --- |
| Comparison | Pseudo-F | R<sup>2</sup> | p-value | $p_B$ | F | p-value | $p_B$ |
| Treatments | 9.4924 | 0.37236 | 0.001* | 0.002* | 0.419 | 0.571 | 1.000 |
| Treatments - Time | 8.6926 | 0.65068 | 0.001* | 0.002* | 4.4403 | 0.021* | 0.041* |

**Table 6.** Summary of results of pairwise post hoc PERMANOVA and Levene’s test for homogeneity of dispersions for tests between all time points within each treatment. (*) means p < 0.05. Bonferroni adjusted p-value (p_B_), k=4.

|  | PERMANOVA |  |  |  | Levene's test |  |  |
| --- | --- | --- | --- | --- | --- | --- | --- |
| Comparison | Pseudo-F | R<sup>2</sup> | p | $p_B$ | F | p | $p_B$ |
| C-38D-[1+2]:C-51D-3 | 6.4619 | 0.48002 | 0.011* | 0.044* | 0.492 | 0.502 | 1.000 |
| C-38D-[1+2]:LP-38D-[1+2] | 11.265 | 0.52975 | 0.008* | 0.032* | 5.0745 | 0.051 | 0.204 |
| C-51D-3:LP-51D-3 | 8.7603 | 0.68653 | 0.1 | 0.4 | 0.5757 | 0.6014 | 1.000 |
| LP-38D-[1+2]:LP-51D-3 | 4.8692 | 0.41024 | 0.011* | 0.044* | 17.507 | 0.001* | 0.004* |

Overall, the bacterial sequences are dominated by a few, common Orders in both treatments, though relative abundance of these groups varies between treatments (Table S4). The primary taxonomic order recovered was Rhodobacterales, and represented nearly 30% of ASVs recovered from each bucket (Fig. 4), but had a significantly lower relative abundance in LP-51D-3 than in other treatment-time categories (Table S5). The Order Cellvibrionales has a significantly higher relative abundance in control buckets at 38 days, but this difference disappears at 51 days. Phycisphaerales relative abundance increased substantially from 38 to 51 days in both treatments, though this change was only significant for the low-pH treatment. Similarly, members of Acanthopleuribacterales are completely absent in buckets at 38 days in both treatments, but appear in low abundance in both treatments at 51 days. Caulobacterales had a higher relative abundance in the low-pH treatment at both time points.

**Figure 4.**
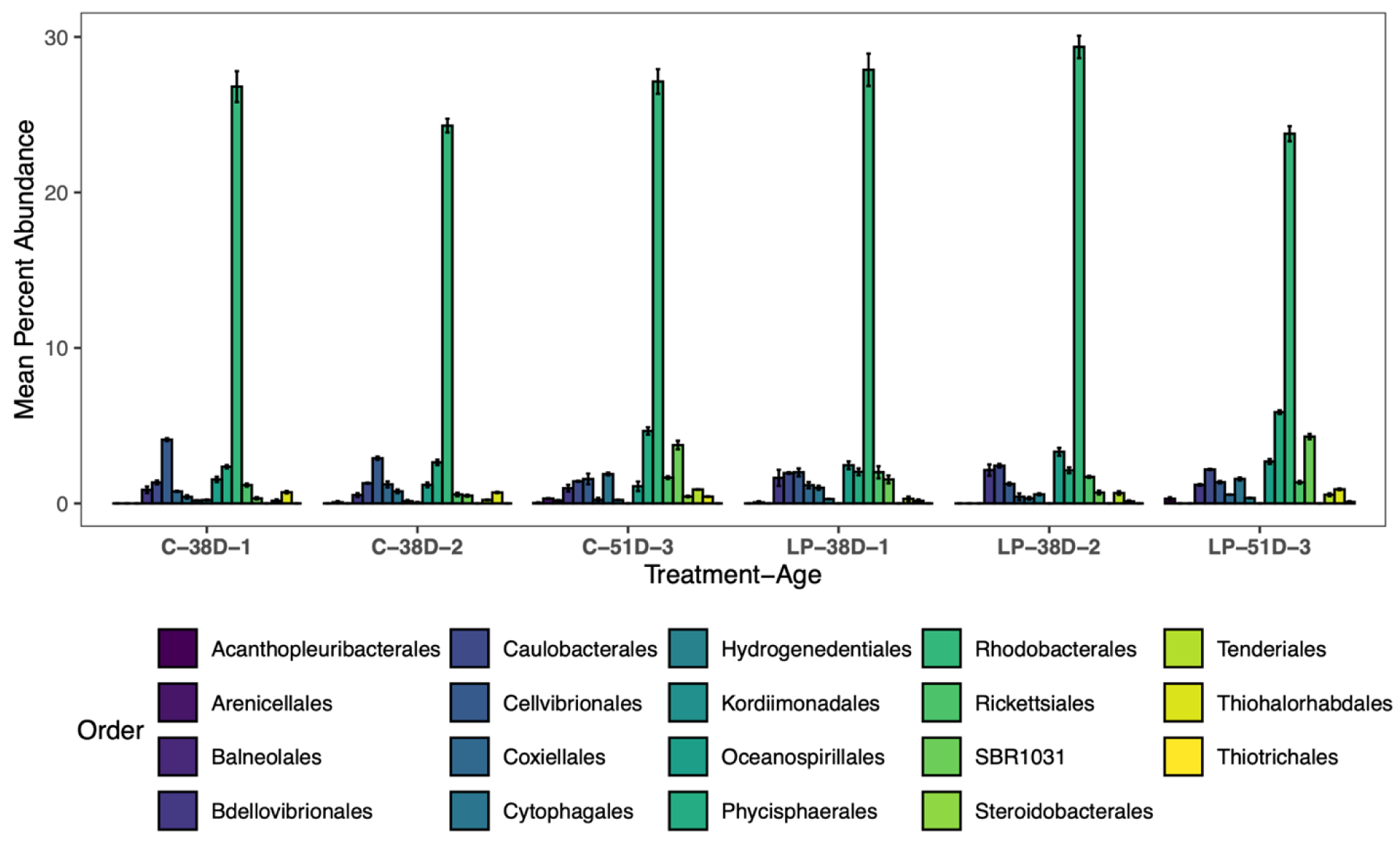
Taxonomic bart chart of most common bacterial Orders. Represents Orders that vary > 0.02% (see Methods) and had significantly different abundances when compared between treatment-time groups.

Changes in microbiome functional abundance were associated with shifts in biological pathways as reconstructed by applying the PICRUST2 program to 16s rRNA data. At Day 38, oyster microbiomes displayed 13 KEGG orthologs that were differentially expressed between the control and low-pH treatments belonging to pathways that affect carbonate precipitation (Table S6). The low-pH treatment had higher function (KO) abundances for orthologs belonging to sulfate-reduction, denitrification, and ammonium oxidation biochemical pathways (Fig. 5). By contrast, oysters from the control displayed higher abundances of functions belonging to photoautotrophy, nitrate reduction, methylotrophic methanogenesis, hydrogenotrophic methanogenesis, and acetoclastic methanogenesis (Fig. 5). Taxonomically, all pathways investigated that are predicted to affect carbonate chemistry were dominated by members of Proteobacteria. Orthologs belonging to sulfate-reduction pathways were also contributed to by members of Planctomycetes and Bacteriodetes, while hydrogenotrophic methanogensis received significant contributions from Chloroflexi (Fig. 5). Generally, these findings suggest several pathways by which microbial taxa could mediate the host environment.

**Figure 5.**
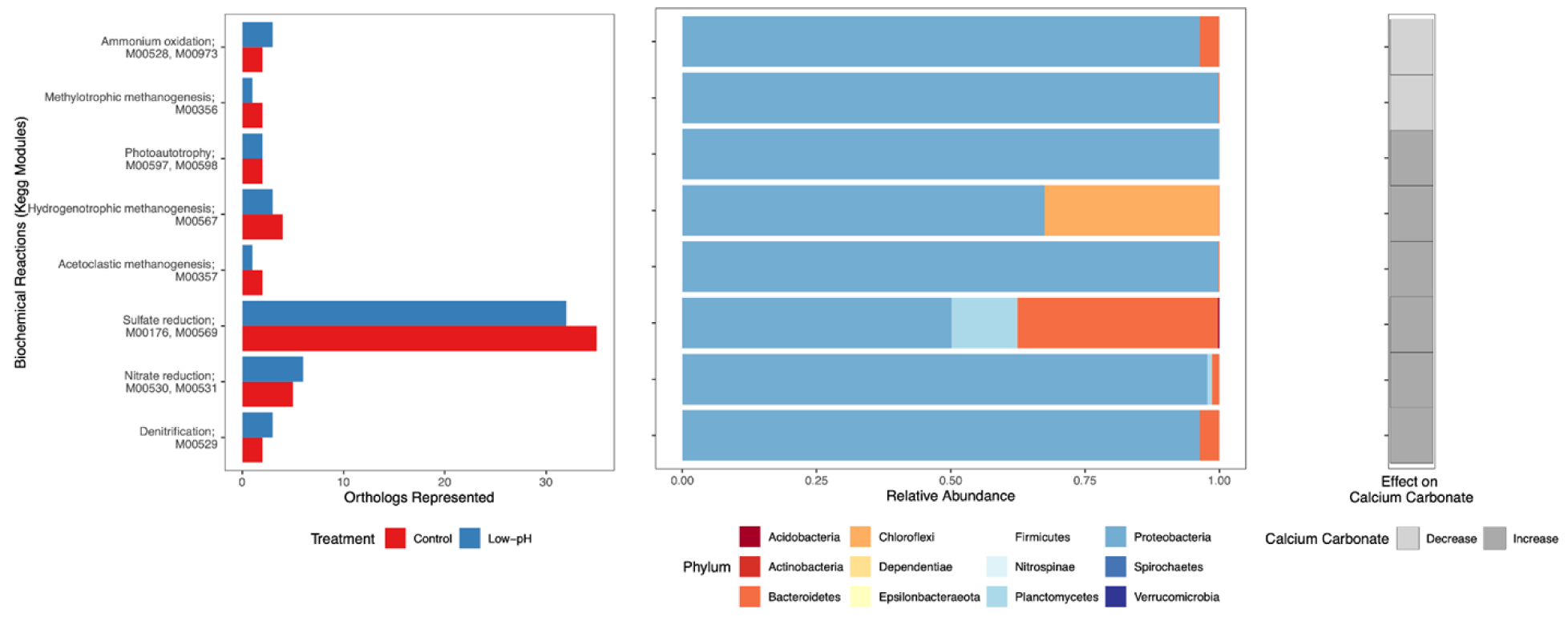
Distribution of KEGG orthology (KO) function abundances and taxonomic composition among microbially mediated biochemical pathways known to affect carbonate chemistry. Data shown belongs to samples collected from 38 day old oysters only. Left: Comparison of KO function abundances among biochemical reactions (KEGG modules). Middle: relative proportions of bacterial Phyla contributing to each pathway. Right: Effect of a given biochemical pathway on carbonate chemistry. Increase indicates that that pathway is likely to increase carbonate saturation state and promote precipitation, whereas decrease indicates that pathway will decrease carbonate saturation and negatively affect carbonate precipitation.

### Differential gene expression analysis

By mapping our RNA-Seq data to the *M.gigas* genome, we generated 39,656 gene models, 36,611 of which matched to gene models in the genome. A total of 3,793 genes were differentially expressed according to cuffdiff, which uses a false discovery adjusted p-value of 0.05. Principal component analysis and heat map clustering of the datasets demonstrates that experimental conditions can be readily distinguished, demonstrating that each experimental condition has a distinct genetic signature (Fig. 6).

**Figure 6.**
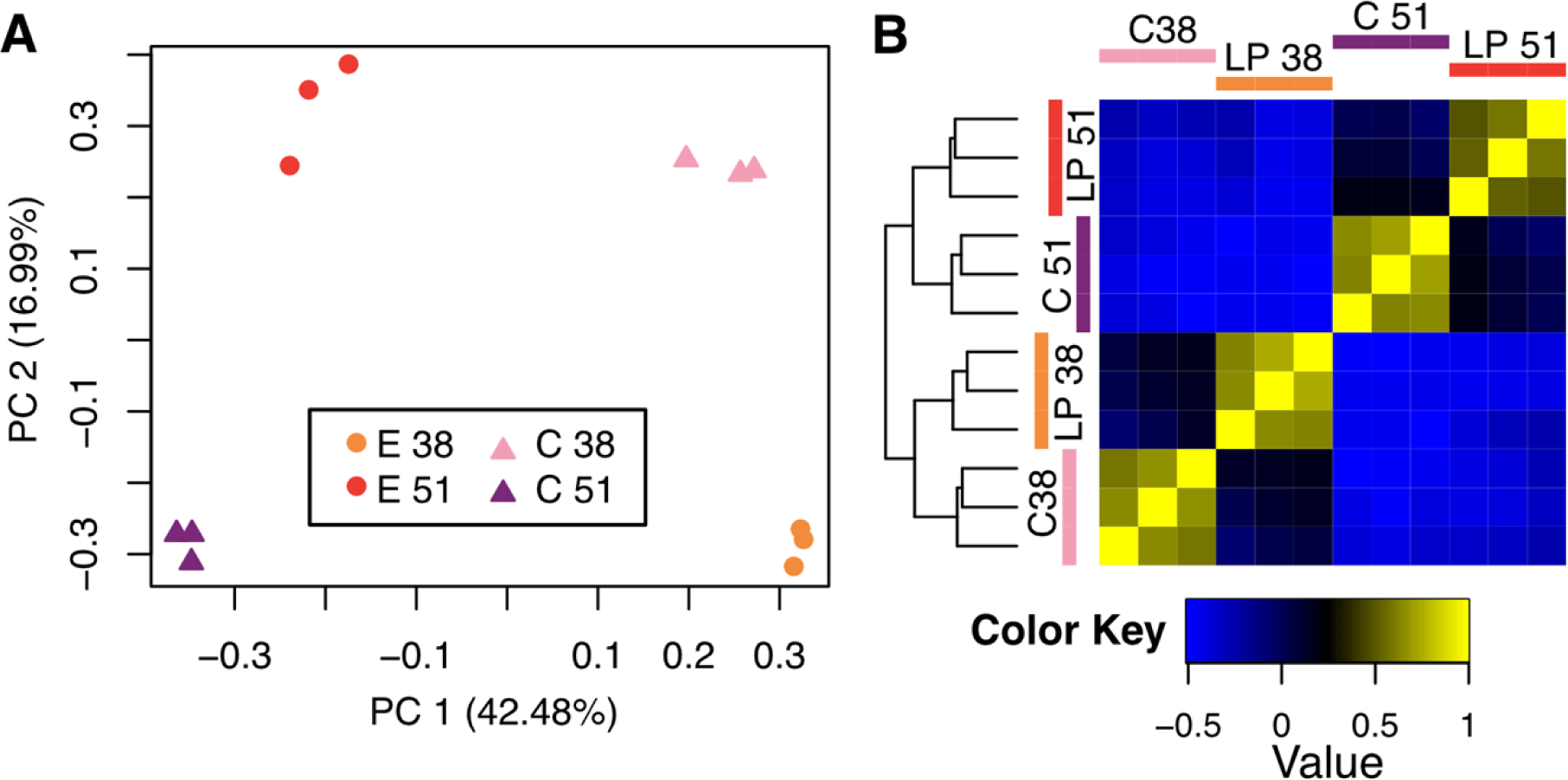
Variation between experimental conditions based on differentially expressed genes. Gene expression data summarized as (A) a principal component analysis plot, and (B) a correlation plot.

We used DAVID functional annotation tools to determine what biological processes were enriched within the RNA-Seq data (Table S7). There were few enriched processes in the control between 38 and 51 days; though a notable enrichment includes copper ion binding / neurotransmitter processes. A different and larger set of enriched processes mark the low-pH condition between 38 and 51 days; notable processes include chitin-binding, calcium ion binding, and collagen / extracellular matrix organization. Comparing the control and low-pH conditions at 38 days, enriched processes included ETS-family proteins, copper ion binding / superoxide dismutases, chitin-binding, and bromodomain-containing proteins/chromatin remodeling. Comparing the control and low-pH conditions at 51 days, enriched processes included zinc-finger / metal binding and heat shock / stress response. This first-order analysis suggests metal ion binding, chitin binding, calcium binding, and heat stress responses were important drivers of shifts in gene expression data.

Digging into the gene annotation, we discovered additional important processes related to biomineralization and microbial interactions. One notable class of genes are carbonic anhydrases, which regulate pH and catalyze the production of calcium carbonate. Five carbonic anhydrases were differentially expressed in our dataset, including two nacrein-like proteins that have been previously identified in the oyster’s shell matrix [49] (Fig. 7). In one case, a nacrein was significantly upregulated in the low-pH condition compared to the control at 38 days (gene ID: LOC105335878), while two carbonic anhydrases– one classified as a nacrein–-were significantly downregulated in the low-pH condition at 51 days (gene IDs: LOC105322241; LOC105327172). Expression patterns for other processes related to biomineralization and microbial interactions are illustrated using heat maps in Fig. 8. This analysis is restricted to genes that were differentially expressed between the control and low-pH conditions at concurrent time points. Genes involved in immune responses were generally downregulated in the low-pH condition, with the exception of three genes (a toll-like receptor, a β-Glucan-binding protein, and an interferon−induced protein) all of which were upregulated in the low-pH condition at 51 days. There was similarly a downregulation of phagosome-related genes in the low-pH condition, as well as downregulation of most chitin binding genes by day 51. The significance of these genes will be considered more in the Discussion, but at a broad level these findings highlight the intricate and varied responses of genes involved in biomineralization and microbial interactions to low-pH conditions.

**Figure 7.**
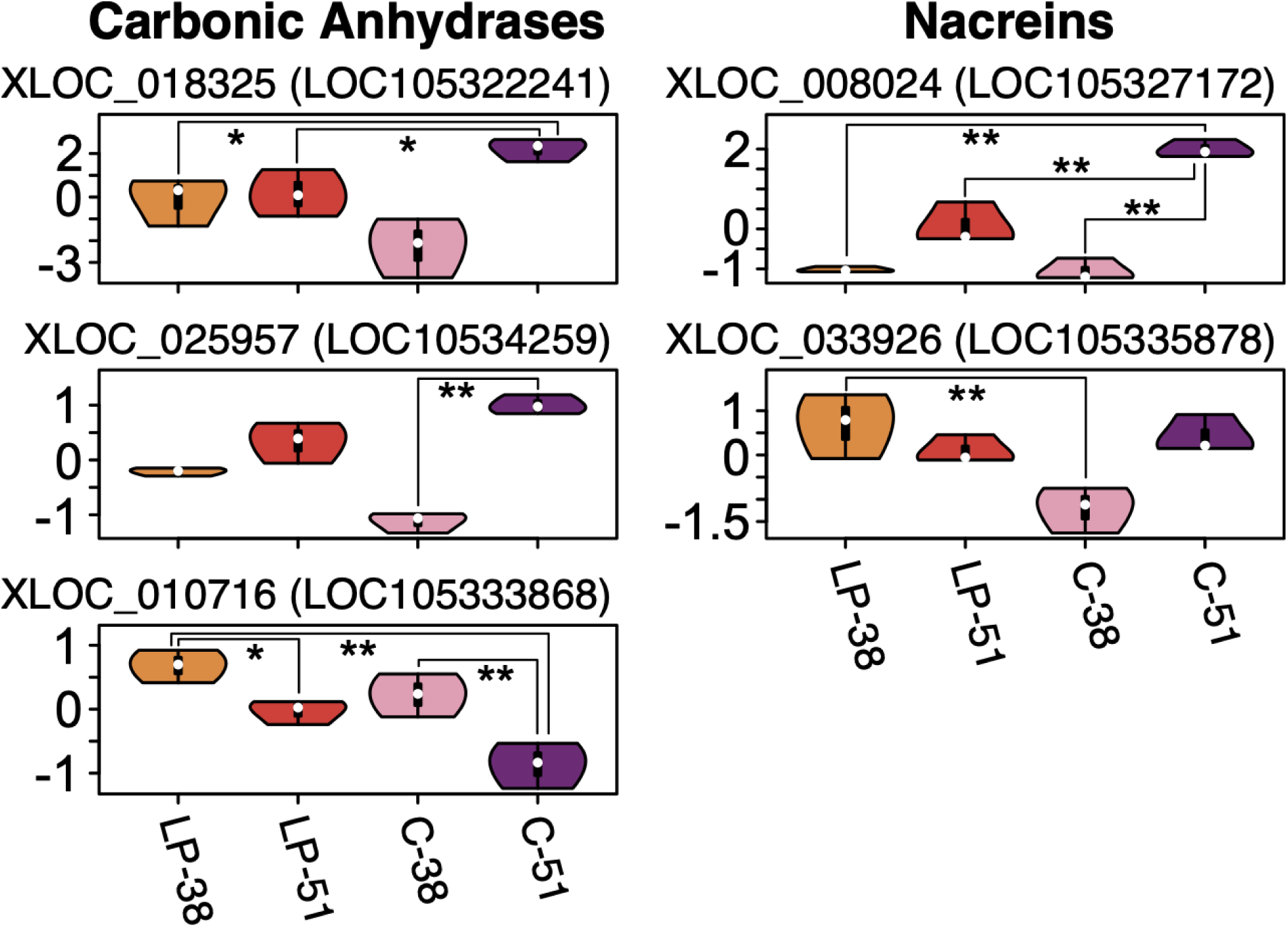
Expression of carbonic anhydrases in our study. Comparisons with a false discovery rate-adjusted p-value of < 0.05 are noted with an asterisk; those with a p-value < 0.01 are noted with two asterisks.

**Figure 8.**
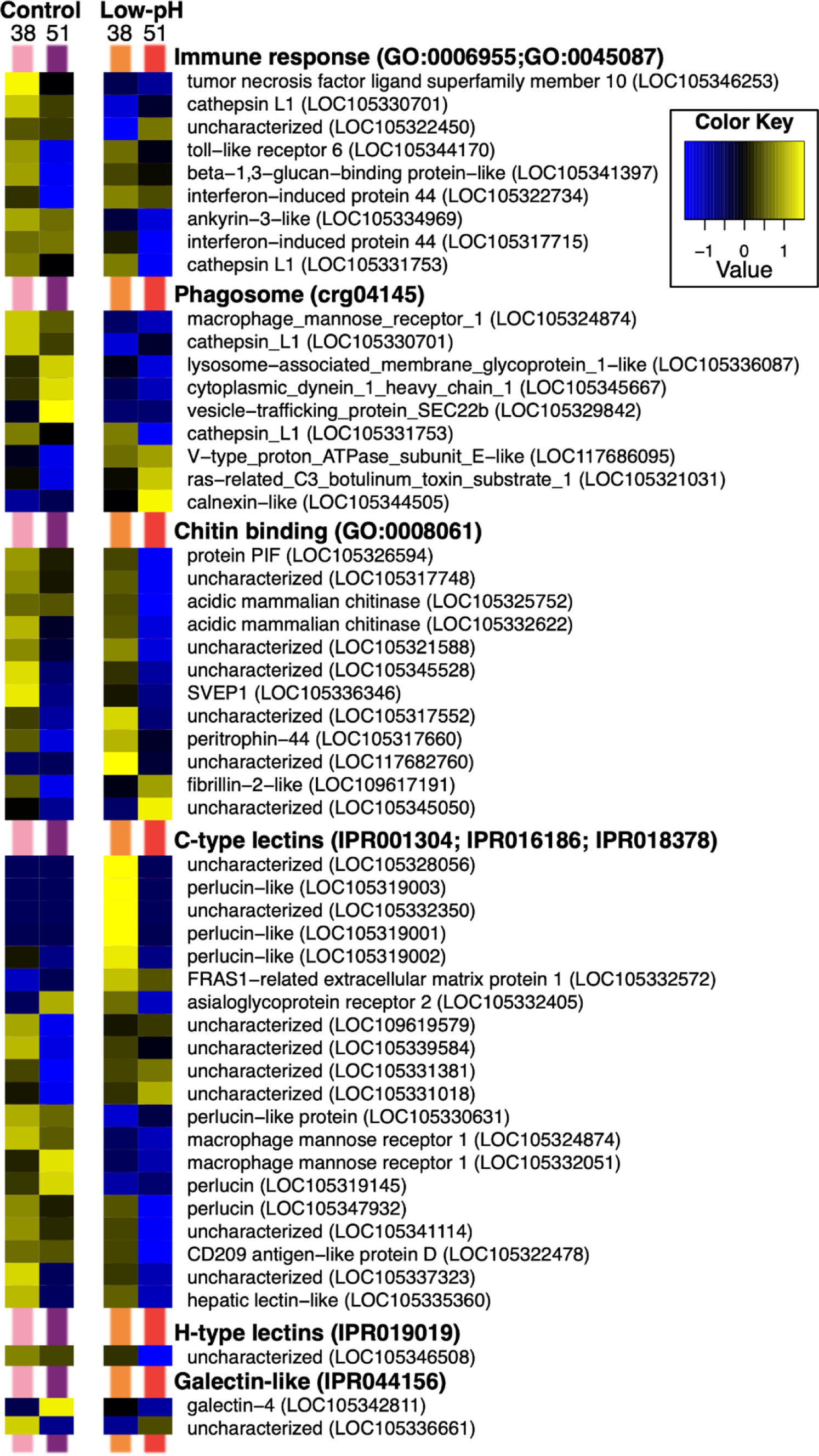
Heat map of important genes in our study. The normalized expression values were log transformed and z-scaled, with values for each gene centered around the mean.

## DISCUSSION

This study demonstrates the interconnected effects of low-pH conditions on shell calcification, microbial communities, and gene expression in juvenile Pacific oysters (*Magallana gigas*). Results reveal that oysters selectively bred for low-pH resistance exhibit increased shell mass under elevated pCO_2_, highlighting the potential of genetic breeding programs in mitigating climate change impacts on the aquaculture industry and marine calcifiers more broadly. This is consistent with previous studies that suggest increased oyster shell growth is connected with adaptation to low-pH seawater ([18]; [21]). However, this trait appears to be time-dependent, with significant shifts in shell growth patterns that correlate with changes in microbial and genetic signatures.

The low-pH treatment applied to juvenile Pacific oysters had a clear effect on microbial community composition (Table 6; Fig. 3), even though it did not uniformly decrease microbial diversity in low-pH samples (Fig. 2). Previous studies have likewise found that while changing ocean pH alters microbial community composition, the diversity and abundance of bacteria are not consistently affected (Krause et al., 2012). This underscores the complexity of microbial responses to environmental stressors, where compositional shifts do not always correspond to changes in overall diversity metrics.

Among microbial biochemical reactions that influence carbonate chemistry, we recovered evidence for shifts in microbial-based ammonium oxidation, methylotrophic methanogenesis, photoautotrophy, hydrogenotrophic methanogenesis, acetoclastic methanogenesis, sulfate reduction, nitrate reduction, and denitrification, as has been found in oyster metagenomic profiles [44]. The increase in sulfate reduction under low-pH conditions was particularly strong, suggesting that acidification may enhance these anaerobic microbial respiration pathways within oysters. Interestingly, we also observed an increase in the denitrification pathway, which is predicted to decrease carbonate saturation state. It is possible that the positive effects of bacterial sulfate reduction create net favorable conditions for carbonate precipitation in low-pH oysters. Taken collectively, this may explain the observation among numerous studies that low-pH conditions affect calcification differently among different oyster taxa. Depending on the microbial community composition within a given oyster, which is influenced by environmental conditions and host species identity [13,14], changing ocean pH conditions may favor certain microbial taxa over time. Depending on the initial microbial source pool, and the selective pressures of environmental change, this could result in a microbial population that increases or decreases carbonate saturation state. Given that low-pH oysters here experienced greater shell growth than their control counterparts, and that we observe a greater abundance of sulfate-reducing functional orthologs, our work also implies a connection between microbial sulfate reduction as a mechanism for improving carbonate chemistry for metazoan precipitation, as has been suggested previously in oysters [11,13,50].

Consistent with our growth and microbiome data, gene expression profiling of *M. gigas* reveals an enrichment of transcripts related to biomineralization and microbial interactions / innate immunity. Previous studies have noted a strong overlap between these two processes in bivalves such as *M. gigas*, as both are regulated by subpopulations of hemocyte cells [51–53]. Lectins, for example, are a diverse group of carbohydrate-binding proteins, some of which are used by hemocytes for antimicrobial recognition [54], while others (notably perlucins) nucleate the growth of calcium carbonate crystals during biomineralization [55]. Three perlucins were upregulated in the low-pH condition at 38 days, but by 51 days the expression of all perlucins went down, which is consistent with an initial burst in shell growth followed by a slowing down in calcification rate. Though further work would be needed to fully characterize the relationship between the expression of perlucins and shell calcification rate at different ontogenetic stages. Other genes also indicate a shift in hemocyte function. For example, when *M. gigas* is infected with the bacteria *Vibrio parahaemolyticus*, hemocytes identify this pathogen using Macrophage Mannose Receptor 1, and the infection is associated with increased expression of lysosome-associated membrane glycoprotein 1, calthepsins, and SVEP-like proteins [56]. In our dataset, all of these genes are downregulated in the low-pH condition, suggesting that decreased pH results in animals relaxing their immune response. The downregulation of cathepsins could specifically indicate the reduction of granulocytes: the cathepsin gene is strongly associated with granulocytes in the closely related Eastern oyster, *C. virginica*, and is upregulated in response to infection by the parasite, *Perkinsus marinus* [57]. Chitinases, which are downregulated in the low-pH condition, are also indicative of *M. gigas* granulocytes, where they play a role in the immune response [58]. Our results add to a growing literature that entangles immunity and biomineralization in *M.gigas*, which could help explain its adaptive response to low-pH waters and other stressful environments. It also suggests there may be a tradeoff to maintaining calcification under low pH that comes at the expense of increased disease susceptibility.

The differential expression of carbonic anhydrases among treatments and time suggests animal responses to the environment to regulate shell production and pH balance in the calcifying fluid. The two nacrein-like carbonic anhydrases from our study have been previously identified in the shell-forming mantle tissue of *M. gigas* [49]. Song et al. found that gene LOC105327172 (called “nacerin protein F1” in the text) is upregulated in juvenile spat with the start of calcite production, and LOC105335878 (protein “F2”) similarly increases in the juvenile stage, with highest expression in the right mantle valve. Our results show that, under low-pH conditions, *M. gigas* had a higher expression of LOC105335878 (“F2”) than the control at 38 days, but a lower expression of LOC105327172 (“F1”) at 51 days compared to control animals. These results suggest that under low-pH conditions, *M. gigas* may prioritize the expression of nacrein “F2” (gene LOC105335878) to enhance calcite production during early development, while the reduced expression of nacrein “F1” (gene LOC105327172) in low-pH oysters compared to control animals at 51 days could indicate a shift in energy allocation away from subsequent juvenile shell maintenance.

Overall, the results of our study are consistent with the “dysbiosis hypothesis” presented in our previous analyses (Banker et al. 2022). In that study, we attempted to manipulate the microbiome of the same *M.gigas* cohort using sodium molybdate. At 38 days, the microbiomes of the control and experimental (molybdate added) populations differed, and the experimental animals had larger shells on average. By 51 days, neither microbiome nor shell size differed as a function of molybdate treatment. We hypothesized that the experimental animals adjusted their microbial communities through shifts in gene expression, in order to restore their microbiome to maintain shell growth. Here we suggest a similar explanation for the convergence with age of shell size, microbiome composition, and oyster gene expression across pH treatments in the present study, though we acknowledge that reduced replication at day 51 could reduce our statistical power. Moreover, we provide evidence that two nacrein-like carbonic anhydrase genes may be mechanistically responsible for the patterns of shell growth presented here. Our oysters were selectively bred for low-Ph tolerance and the evidence suggests that these organisms could compensate for ocean acidification by leveraging genetic and microbial adaptations. Assessing this ability for natural stocks or other populations will be required to confirm whether this ability is widespread, or restricted to this particular line. Future studies should investigate the long-term stability of these traits, their consistency across oyster strains, and their ecological implications. Understanding the interplay between microbiomes, gene expression, and environmental variables will be crucial for developing sustainable aquaculture practices and ensuring the future health of marine bivalve populations.

## ACKNOWLEDGEMENTS

We gratefully acknowledge the Hog Island Oyster Company for supplying us with broodstock oysters for this experiment. Thanks also to Joe Newman and the rest of the Aquatic Resources Group at the Bodega Marine Laboratory.

## COMPETING INTERESTS

The authors have no competing interests to report.

## FUNDING

Funding was provided by the University of California Davis Microbiome Special Research Program (to R.M.W.B, J.J.S., and D.A.G.) as well as NSF award 2044871 (D.A.G.).

## DATA AND RESOURCE AVAILABILITY

The raw data from the genetic analyses have been submitted to NCBI under bioproject PRJNA762081. Images of shell data are available on Harvard Dataverse, https://doi.org/10.7910/DVN/QOLCCA. The code as well as intermediate files are preserved on GitHub: https://github.com/DavidGoldLab/2025_Oyster_RNA-Seq and https://github.com/Roxanne-Banker/Oyster_OA

## Notes

### Competing Interest Statement

The authors have declared no competing interest.

